# Haplotype-resolved assembly of a tetraploid potato genome using long reads and low-depth offspring data

**DOI:** 10.1101/2022.05.10.491293

**Authors:** Rebecca Serra Mari, Sven Schrinner, Richard Finkers, Paul Arens, Maximilian H.-W. Schmidt, Björn Usadel, Gunnar W. Klau, Tobias Marschall

**Affiliations:** Institute for Medical Biometry and Bioinformatics, Medical Faculty, Heinrich Heine University, Düsseldorf, Germany; Algorithmic Bioinformatics, Heinrich Heine University Düsseldorf, Germany; Gennovation B.V., Agro Business Park 10, 6708 PW, Wageningen, The Netherlands; Plant Breeding, Wageningen University & Research, The Netherlands; Cluster of Excellence on Plant Sciences (CEPLAS), Heinrich Heine University Düsseldorf, Germany; Forschungszentrum Jülich, Institute of Bio and Geosciences, Bioinformatics (IBG-4), Germany; Bioeconomy Science Center, c/o Forschungszentrum Jülich, Germany; Biological Data Science, Heinrich Heine University Düsseldorf, Germany

## Abstract

Potato is one of the world’s major staple crops and like many important crop plants it has a polyploid genome. Polyploid haplotype assembly poses a major computational challenge, hindering the use of genomic data in breeding strategies. Here, we introduce a novel strategy for the assembly of polyploid genomes and present an assembly of the autotetraploid potato cultivar Altus. Our method uses low-depth sequencing data from an offspring population, which is available in many plant breeding settings, to achieve chromosomal clustering and haplotype phasing directly on the assembly graph. This involves a novel strategy for the analysis of *k*-mers unique to specific graph nodes. Our approach generates assemblies of individual chromosomes with phased haplotig N50 values of up to 13 Mb and haplotig lengths of up to 31 Mb. This major advance provides high-quality assemblies with haplotype-specific sequence resolution of whole chromosome arms and can be applied in common breeding scenarios where collections of offspring are available.

## MAIN

Polyploidy is common in plant genomes and two forms are recognized. Allopolyploids arise from interspecific or intergeneric hybridization events, and the difference between subgenomes is usually sufficient to assemble them like diploids. This has been demonstrated for rapeseed, wheat and strawberry, among others (Kyriakidou et al. 2018). In contrast, autopolyploids arise from genome duplications, and the presence of multiple sets of the same homologous chromosomes means that haplotype-resolved sequence assemblies are much more challenging. One example is potato (*Solanum tuberosum*), most cultivars of which are autotetraploid (Petek et al. 2020). Potato is a vital food crop in many developing countries (Devaux, Kromann, and Ortiz 2014), and the global production volume exceeds 300 million tons per year (Birch et al. 2012). Because of this agronomic value, efforts to assemble potato genomes are of crucial importance.

The haplotype-resolved assembly of diploid genomes has been progressively refined, and accurate results are now possible as we have shown previously (Ebert et al. 2021; Porubsky et al. 2021). In contrast, computational methods for polyploid haplotype assembly rarely lead to satisfying results, particularly for autotetraploids. Reference-based approaches for haplotype phasing in polyploid species align reads to an existing reference sequence but are often inaccurate (Motazedi et al. 2018). Especially in the presence of structural variation, reference-based approaches in general have severe limitations (Porubsky et al. 2021). For potato haplotype phasing, two reference genomes are currently used: the synthetic double monoploid potato clone DM1–3 516 R44 (Pham et al. 2020) and Solyntus, which is based on a diploid potato cultivar (van Lieshout et al. 2020). Reference-based algorithms for polyploid haplotype phasing include HapTree (Berger et al. 2014) and H-PoP (Xie et al. 2016). Other methods target selected genomic regions to resolve haplotypes locally, for example using integer linear programming (Siragusa et al. 2019). We previously developed WhatsHap polyphase, which was an improvement over contemporaneous methods but still relied on a reference genome (Schrinner et al. 2020).

The *de novo* assembly of polyploid genomes without a reference is still an emerging strategy. Recently proposed workflows involve the building of a “squashed” assembly with no or limited haplotype resolution at first, and using this as the basis for haplotype phasing. Even long sequencing reads are generally insufficient for long-range phasing, and auxiliary data types are required. One example is single pollen cell sequencing (Zhang et al. 2021), which was recently used for comprehensive haplotype reconstruction in autopolyploid potato (Sun et al. 2022). Here, we propose an alternative method in which PacBio HiFi reads of the potato cultivar Altus are combined with cost-effective low-coverage short-read sequences from multiple offspring samples. Accordingly, we generated PacBio HiFi reads (96× coverage) and created an initial assembly using hifiasm (Cheng et al. 2021). We assembled the individual haplotypes from the resulting assembly graph using sequencing data from 193 offspring of two potato cultivars (Altus and Colomba) at low coverage (∼1.5× per haplotype) combined with a novel approach based on *k*-mers to identify the four haplotypes. Our assembly mapped well to the latest version of the monoploid DM1–3 516 R44 reference (DMv6.1) and yielded haplotype-resolved assemblies of individual chromosomes with phased haplotype block lengths of up to 31 Mb, phased contig N50 values of up to 13 Mb, and a genome-wide phased contig N50 value of 7.2 Mb. Our approach also allows the detection and correction of assembly errors in the assembly graph as well as in previously published references.

## RESULTS

### Overall assembly strategy

A high-level overview of our workflow is shown in Fig. 1. Starting with PacBio HiFi reads derived from the Altus genome (Fig. 1a), we built an assembly graph using hifiasm, resulting in a partially haplotype-resolved graph with bubble-like structures representing the different haplotypes (Fig. 1b). For each so-called unitig in the assembly graph, we detected unique *k*-mers (Fig. 1b). We then estimated the dosage of each unitig, defined as the number of haplotypes to which each unitig contributes (Fig. 1c). In the next step, we counted the formerly detected unique *k*-mers in the Illumina reads for each of the 193 offspring samples (Fig. 1d). Each unitig is thus represented by a *k*-mer count pattern consisting of 193 values. Nodes with similar count patterns, implying the inheritance of a node by the same subset of offspring samples, are therefore likely to be part of the same haplotype. We then made use of the *k*-mer count patterns to perform an initial clustering of the nodes into chromosomes (Fig. 1e). The clustering procedure was followed by a step to determine the four haplotypes among nodes with dosage 1 (Fig. 1f), and another step to add nodes with higher dosages (Fig. 1g). Ultimately, this yielded a set of four haplotype clusters for each chromosome (Fig. 1h). We completed the assembly by finding graph traversals through the clustered assembly graph and thereby assembling haplotype blocks (haplotigs).

**Fig. 1:**
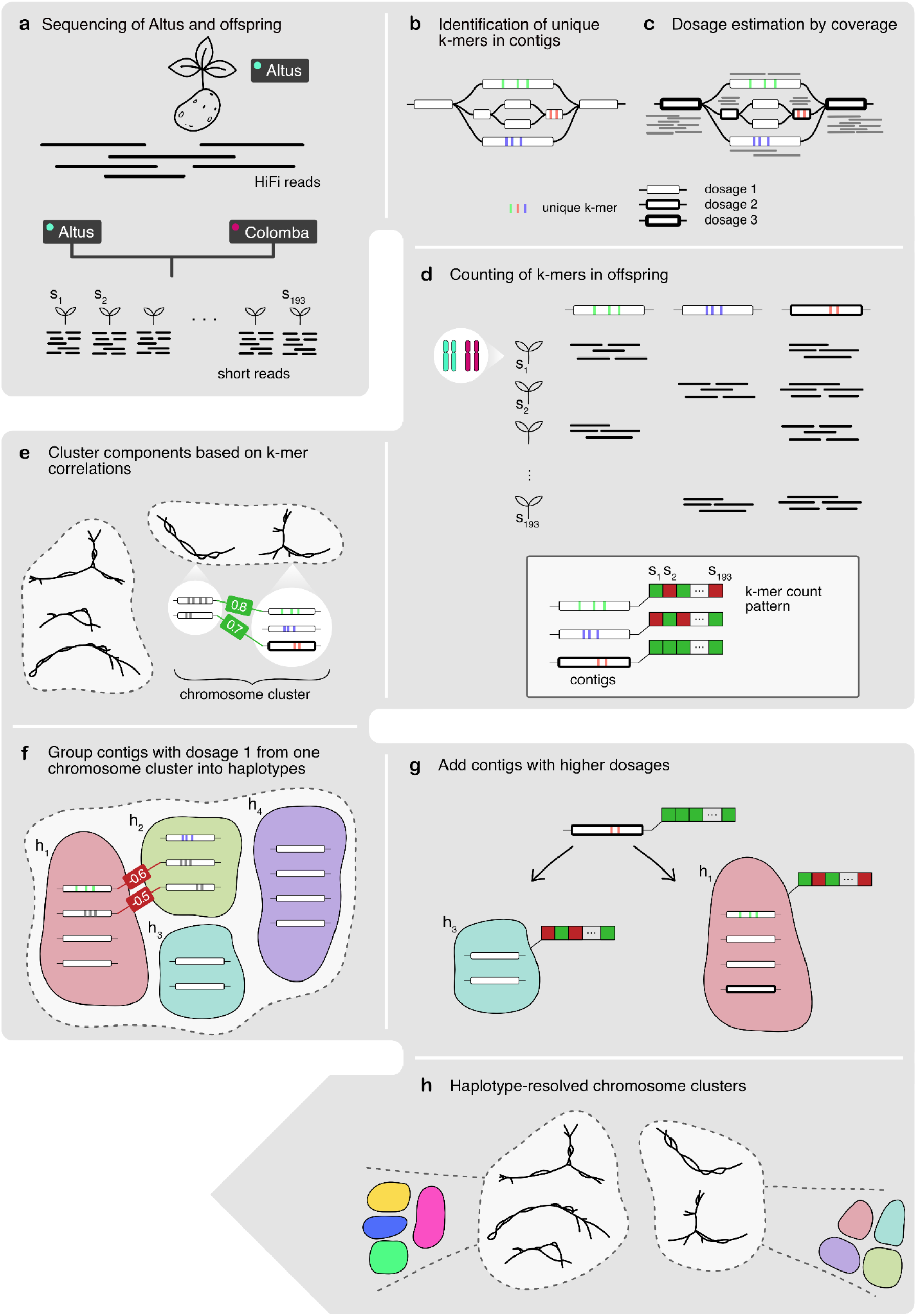
Overview of the workflow. **a**. The Altus genome was sequenced using PacBio HiFi technology, whereas the 193 genomes of the cross Altus × Colomba were sequenced on the Illumina platform. **b**. We used hifiasm to assemble the Altus HiFi reads into an assembly graph. For each contig in the graph, unique *k*-mers were detected (denoted by the colored bars). **c**. The HiFi reads were aligned to the contigs and the mapping depth was used to estimate dosages (1 to 4) for each contig. The different dosages are denoted by the thickness of the contig line (thicker outlines mean higher dosage). **d**. The unique *k*-mers were counted in the short reads of the offspring samples in order to compose a count pattern for each contig. **e**. For all nodes from the assembly graph components, the pairwise correlation of *k*-mer count patterns was computed and components were clustered to represent chromosomes. **f**. In each chromosome cluster, the nodes with estimated dosage 1 were first clustered into the four haplotypes, again based on pairwise correlations. **g**. The contigs with dosages > 1 were added to the clusters that contain most matching nodes in terms of *k*-mer count pattern correlations. **h**. This process resulted in chromosome clusters that contain subclusters for each haplotype.

### Initial assembly

We first sequenced the Altus genome using PacBio HiFi technology to produce highly accurate long reads with an average coverage of 24× per haplotype (73.7 Gb in total). We also acquired Illumina short-read sequencing data representing 193 offspring of the cross between Altus and another cultivar (Colomba). The data consisted of 2 × 150 bp paired-end reads with an average coverage of 1.5× per haplotype.

We assembled the HiFi reads using hifiasm v0.13, which outputs an assembly graph that contains all the assembled, unprocessed (raw) unitigs, which partially resolve the four haplotypes. Variation is represented by bubble structures in the graph, where a unitig branches into two or more other unitigs.

The initial graph consisted of 20,216 nodes (unitigs), 26,566 edges and contained 2798 Mb of sequence data. The N50 value of the unitigs was 1.34 Mb. The nodes of the unitig graph (Supplementary Fig. 1) within the 10 largest connected components covered 91–190 Mb each (1.27 Gb in total), 11 further components covered 45–66 Mb each (555.2 Mb in total) and a set of smaller components covered 20–32 Mb each (249.1 Mb in total). Additionally, 699 unitigs were not connected to any other node. In summary, the initial raw unitig graph provided a certain degree of haplotype resolution, indicated by the total amount of sequence data (3.8× the size of the DMv6.1 reference genome), but did not provide longer-range phasing at many loci, indicated by the substantial number of nodes shorter than 50 kb (Fig. 2a).

**Fig. 2:**
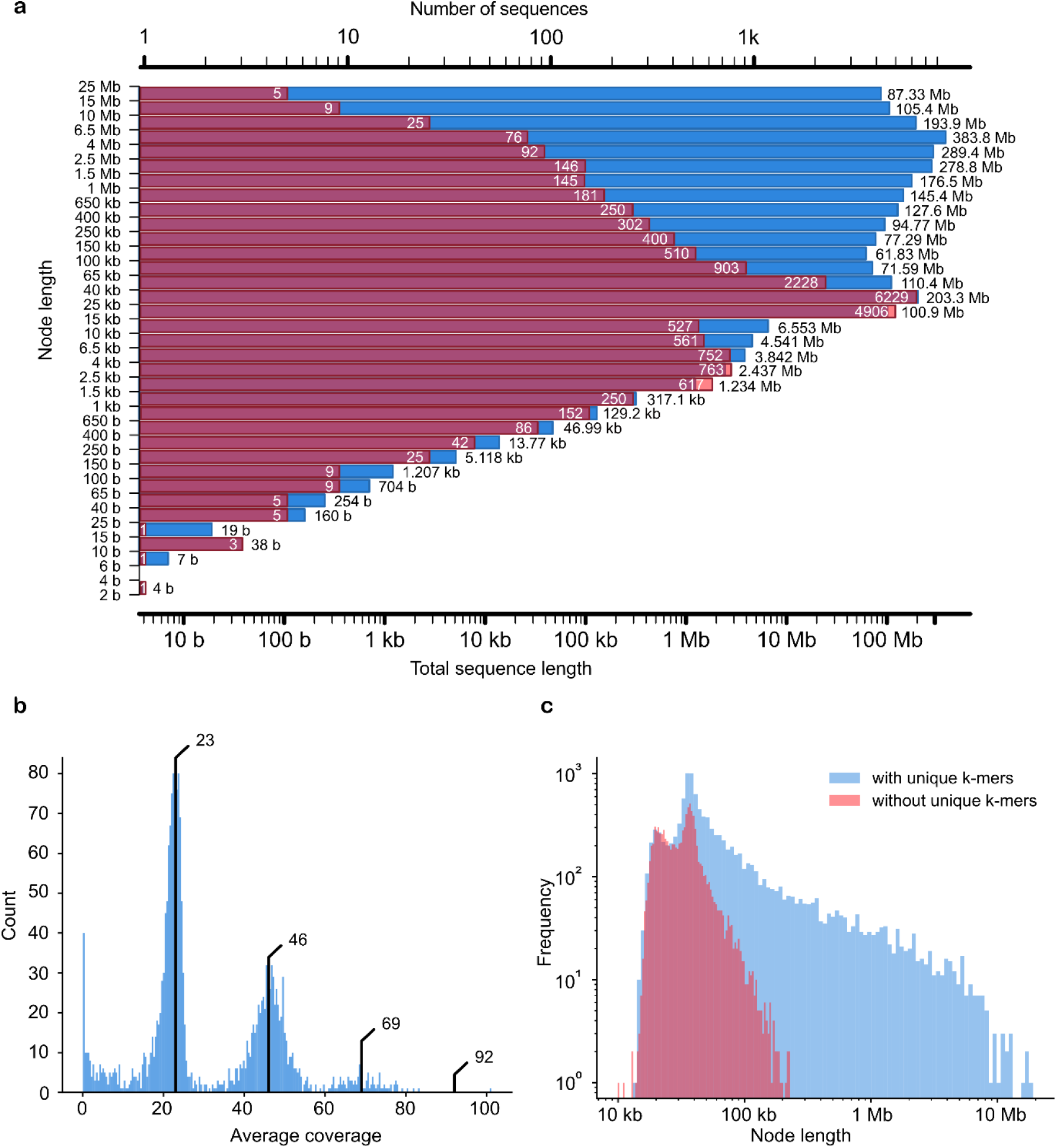
Initial assembly. **a**. Distribution of node lengths of the initial assembly graph. Red represents the count of each binned contig length (the peak is 25–40 kb). Blue represents the aggregate length of a contig bin, measured in bases. The two visible peaks show that the total sequence of contigs between 25 and 40 kb is on par with the sequence taken up by those between 4.0 and 6.5 Mb. **b**. Dosage distribution of contigs, excluding those with a unique sequence < 100 kb. The proportion of sequence that is covered by contigs at least 100 kb in length is 80%. The dosage peaks are marked by black bars (approximate coverage values of 23, 46 and 69). The peak for dosage 4 would be ∼92. **c**. Length distribution of contigs with unique *k*-mers compared to contigs without unique *k*-mers.

### Dosage analysis

For each unitig, we estimated the dosage (number of haplotypes represented), which for a tetraploid genome can be any value from the set {1, 2, 3, 4}. This was achieved by analysing the coverage of reads aligned to the unitigs. First, we aligned all Altus HiFi reads to the graph unitigs using minimap2 (Li 2018) and filtered out all alignments with a mapping quality below 60. Using the remaining alignments, we computed the sequencing depth at each base position. Given that hifiasm graphs usually contain overlaps, we computed the intervals of non-overlapping sequences per node (the region of each node that is not part of any overlap with its neighbouring nodes) and only computed the depth in these unique regions, leading to an average depth per node. Nodes with a non-overlapping sequence 100 kb or longer (Fig. 2b) covered ∼80% of the total sequence in the graph. Three peaks were observed, representing approximate coverage values of 23, 46 and 69, consistent with dosages of 1, 2, and 3. A fourth peak (∼92) was missing for the long contigs (Fig. 2b) and barely visible for all contigs (Supplementary Fig. 2). This may indicate the existence of only a few homozygous regions and the complete absence of long homozygous stretches exceeding 100 kb.

For 6212 contigs, the sequence consisted solely of overlaps to both neighbouring nodes. Given the absence of a unique region, we therefore omitted these contigs from the computation of coverage. Of the 8290 nodes with a depth value above zero, 72.77% were labelled as dosage one, 15.01% as dosage two, 7.95% as dosage three, and 2.97% as dosage four. The remaining 1.3% of the contigs exceeded dosage four and are presumed to represent repetitive regions.

### Analysis of k-mers

In the next step, we counted all possible *k*-mers (fragments of length *k*, in our case *k* = 71) within the unitigs. We then identified those appearing exactly once in the entire graph and reported this set as unique *k*-mers. Approaches based on unique *k*-mers have facilitated the haplotype-resolved assembly of diploid parent-offspring trios (Koren et al. 2018) and challenging regions of human chromosome 8, such as the centromere (Logsdon et al. 2021). In the latter example, the authors created a library of singly unique nucleotide *k*-mers (SUNKs) to barcode long reads and assemble them into scaffolds especially in complex regions. Here, we have developed a novel approach to phase the assembly graph of a parent genome from a polyploid offspring panel. For each unitig, we used the corresponding set of unique *k*-mers as an identifier for the node, making sure the *k*-mers are unique for the Altus genome by disregarding those also found in the Colomba genome. The *k*-mer counting process is based on the Jellyfish API (Marçais and Kingsford 2011).

The resulting set of unique *k*-mers was counted in the 193 offspring samples. Given that each of the tetraploid offspring inherits two haplotypes from Altus and two from Colomba, we inferred the number of inherited copies of a unitig by assessing the abundance of unique *k*-mers for that unitig. Based on the unitig dosage in the parental Altus genome, there are different possible dosages in the offspring. For dosage 1 in the parent, the *k*-mer representing the unitig can be absent or present in an offspring genome, whereas for parental dosage 2, the *k*-mer can be absent, present once, or present twice in the offspring. For parental dosage 3, dosages of 1 and 2 are possible in the offspring, and for parental dosage 4, both inherited haplotypes must arise from this unitig.

Based on the above, we can denote unitigs with unique *k*-mers as *phase informative* and those without unique *k*-mers as *phase uninformative*. The analysis of node lengths for the sets of phase informative and uninformative nodes is shown in Fig. 2c. As anticipated, the uninformative unitigs were generally the shorter ones. Among the complete set of 20,216 nodes, we found that 10,784 (53.34%) were phase informative. Recall that 6212 contigs did not have a unique region due to overlaps, so that unique *k*-mers cannot be present in these nodes. The length of the sequence covered by informative nodes in relation to the sequence covered by all nodes was 88.15% (2.466 of 2.798 Gb), showing that phase uninformative nodes tend to be shorter than phase informative nodes. Specifically, the average node length in the set of phase informative nodes was 228.7 kb (N50 = 1.89 Mb) whereas the average for uninformative nodes was 35.1 kb (N50 = 37 kb). The longest unitig without a unique *k*-mer was 237 kb, compared to 19.11 Mb for the longest informative unitig. Thus, despite the relatively high number of phase uninformative nodes, most of the sequence (88.15%) was generally amenable to offspring-based phasing using our technique.

### Correlation analysis

For our correlation-based approach, we assumed that contigs from the same haplotype have similar *k*-mer count patterns because they occur on the same subset of offspring samples. Therefore, we computed Spearman correlation coefficients (ρ) between the *k*-mer count patterns and analysed the distribution of correlations throughout the assembly graph. Genomic loci that are at close distance (and thus, tightly linked) are likely to be transmitted together to progeny, and accordingly, the offspring-based haplotype signal gets weaker with increasing distance due to recombination. In line with this expectation, we observed a strong correlation for contigs at distances < 10 Mb and decreasing correlation for greater distances (Fig. 3a). For distances below 10 Mb, we observed a bimodal distribution indicative of contig pairs on the same and on different haplotypes.

**Fig. 3:**
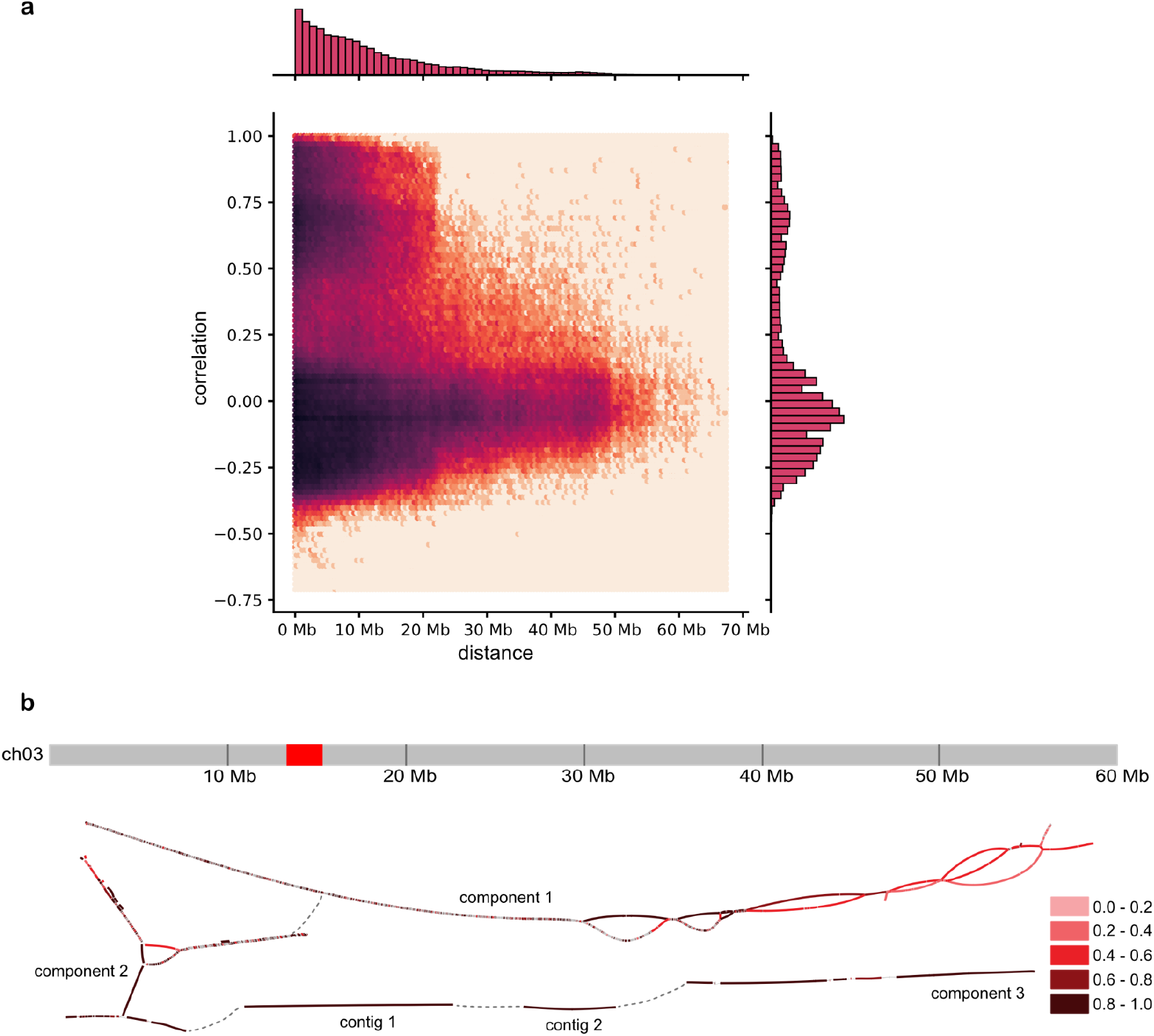
Correlation analysis. **a**. The correlation of all node pairs (nodes with dosage 1) in the 20 largest connected components as a function of the distance between nodes (in megabases). **b**. Reconstruction of the structure of chromosome 3 based on high correlation coefficients between nodes. Chromosome 3 is shown above, with the red block labelling the centromere as reported in the DMv6.1 annotation. The initial assembly consisted of three connected components and two additional contigs, which were manually placed at their approximate genomic location along the *x*-axis as determined by mapping the unitigs to DMv6.1 (the darker the colour of a contig, the higher the maximum correlation to any other contig beyond its component). Contig pairs with the highest correlation (here denoted by the darkest colour, representing a correlation coefficient of 0.8–1.0) could then be connected, revealing a more complete structure of the haplotype-resolved chromosome. The connected node pairs are marked by the dotted grey line.

Based on high correlation values, we were able to reconstruct areas from the graph that were unconnected in the initial assembly, such as broken bubble structures and unconnected fragments. A representative reconstruction of chromosome 3 is shown in Fig. 3b. In the initial assembly, this chromosome consisted of three connected components and two longer unconnected contigs. By connecting contig pairs with very high positive correlation coefficients (ρ = 0.8–1.0), we were able to reconstruct the phased structure of the chromosome and to order the components and contigs accordingly.

### Graph traversal and final haplotype assembly

We assigned each unitig to a haplotype based on our novel clustering procedure, which is described in more detail in the Online Methods. We extended the clustering beyond the graph components and thereby matched the components belonging to the same chromosome. This resulted in 12 pseudo-chromosomes, each consisting of four clusters of haplotagged unitigs. Additionally, we connected the resulting clustered contigs as far as possible by finding graph traversals within the assembly graph, yielding blocks corresponding to the four haplotypes, which we describe as haplotigs. The longest haplotig per chromosome ranged from 13.84 Mb on chromosome 10 to 31.99 Mb on chromosome 11. The haplotig N50 value was ≤ 12 Mb and the total N50 value was 7.17 Mb. The full dataset is presented in Table 1.

**Table 1:** Comparative length of the reference genome, the phased assembly after clustering into haplotypes, and the phased assembly after constructing the final haplotigs. The phased length is defined as the sum of the contig lengths contained in the four haplotypes for each chromosome. The N50 value is computed with four times the reference length as the underlying genome size.

| Chromosome | Length of DMv6.1 (Mb) | N50 (Mb) | Longest haplotig (Mb) | Sum of haplotigs (Mb) |
| --- | --- | --- | --- | --- |
| 01 | 88.59 | 5.45 | 25.62 | 336.93 |
| 02 | 46.10 | 13.57 | 24.22 | 243.73 |
| 03 | 60.71 | 7.46 | 27.81 | 221.22 |
| 04 | 69.24 | 4.96 | 15.19 | 283.36 |
| 05 | 55.60 | 10.44 | 19.85 | 218.84 |
| 06 | 59.09 | 12.19 | 31.37 | 234.13 |
| 07 | 57.64 | 9.47 | 24.22 | 174.88 |
| 08 | 59.23 | 4.54 | 21.75 | 222.27 |
| 09 | 67.60 | 6.99 | 17.59 | 232.99 |
| 10 | 61.04 | 3.46 | 13.84 | 243.39 |
| 11 | 46.78 | 11.93 | 31.99 | 171.28 |
| 12 | 59.67 | 8.74 | 20.75 | 218.84 |
| Total | 731.29 | 7.17 | 31.99 | 2801.26 |

We compared the assembled pseudo-chromosomes to the latest version of the monoploid reference, DMv6.1 (Pham et al. 2020). To compute the N50 measures, we estimated the tetraploid genome size by using fourfold the length of DMv6.1. The cumulative size comparison of each chromosome based on our assembled pseudo-chromosomes and DMv6.1 is provided in Table 1. The size of the individual phased chromosome was 3.5–4 times as large as the reference, and the total phased length was ∼3.8 times as large, consistent with structural variation and sequence loss on some of the haplotypes as previously observed for other cultivars (Sun et al. 2022).

For the comparison of the Altus assembly and DMv6.1, we mapped the resulting clusters to the reference using minimap2. The corresponding mapping intervals (Fig. 4) indicated that all chromosomes in the assembly were nearly complete, and no large gaps were detected. In all chromosomes, the assembly consisted entirely of contigs from one single cluster, supporting the robustness of our clustering process.

**Fig. 4:**
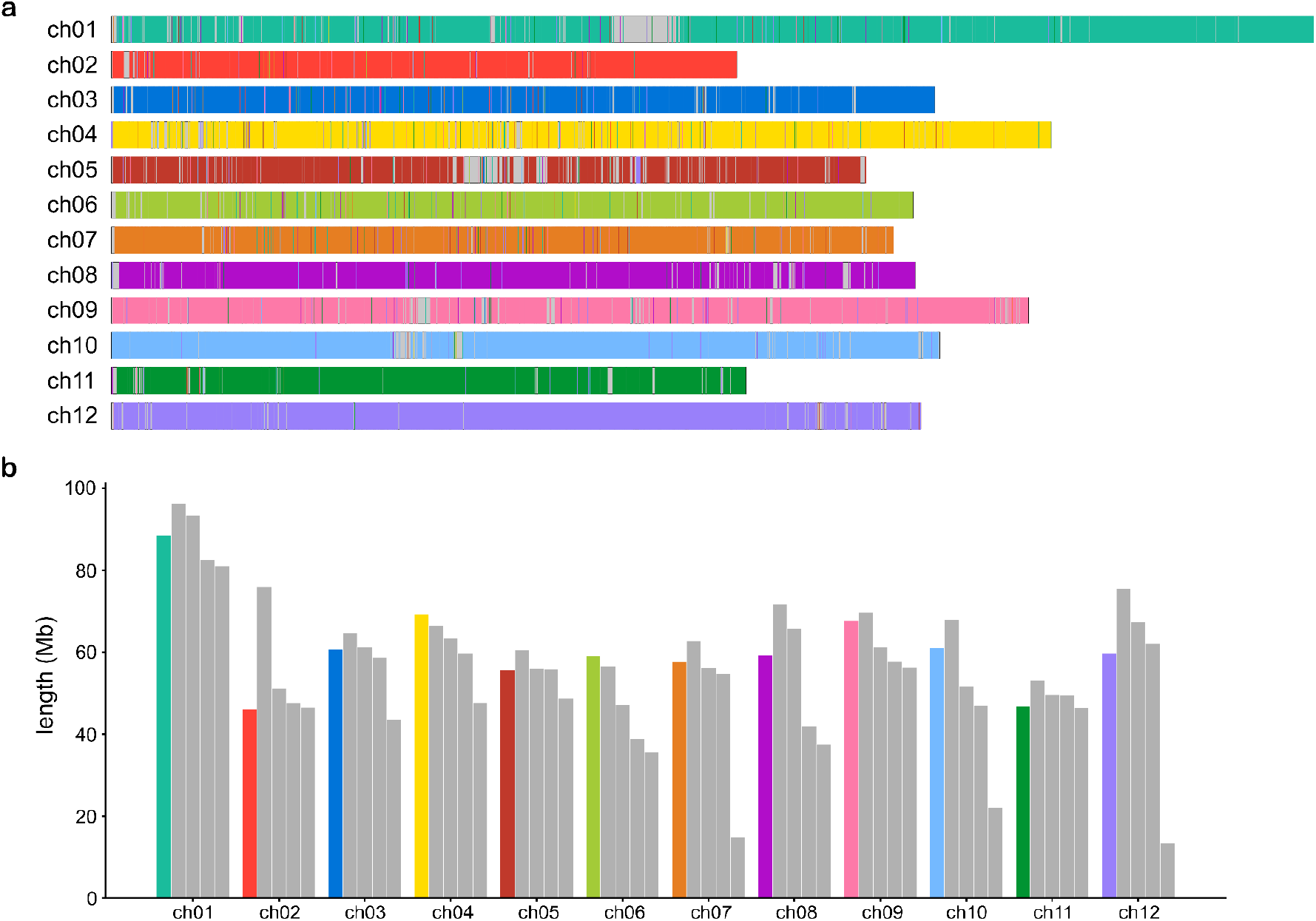
Clustering results. **a**. The contigs of each chromosome cluster are mapped to the reference sequence DMv6.1, and the mapped interval is coloured accordingly. A different colour is used for each cluster. Ideally, one chromosome contains a single colour. **b**. Length comparison of the four haplotypes (grey bars) compared to the reference (coloured bars). The length is computed as the sum of the contig lengths for all contigs in a haplotype cluster.

### Comparison of earlier reference assemblies to reveal structural differences

The correlation signal underlying our chromosome clustering approach was used to detect structural differences between our assembly graphs and previous reference assemblies. Such differences can indicate assembly errors in either of the two assemblies, as well as structural differences in all or some haplotypes. When comparing the initial assembly graph (the hifiasm output) to the DMv6.1 reference, we detected two sets of nodes present on the same component of the graph that mapped to different chromosomes in DMv6.1 (Supplementary Fig. 3). For two contig sets on separate chromosomes, we would expect to see little to no correlation between node pairs from the two sets. Indeed, for the two sets in question, the correlation distribution was very similar in shape to the correlation between one of the sets and a comparison set from a different chromosome (Supplementary Fig. 3a). This probably indicates a false join in the hifiasm graph, which we corrected by manual curation. In this way, correlation analysis provides an opportunity to detect and correct residual assembly errors.

We then compared our assembly graphs to the diploid reference Solyntus (van Lieshout et al. 2020) and found a number of larger structural differences (Supplementary Fig. 4). One example can be found in chromosome 8, where two regions are assembled from contigs that belong to the same clusters as chromosome 7 and chromosome 1, respectively. To investigate whether this was a clustering artefact, an error in the Solyntus assembly, or a true structural difference, we mapped the connected components from the graph representing chromosome 8 individually to the Solyntus reference and identified one component that contained a large fragment of chromosome 1 but also the inserted region on chromosome 8 (Supplementary Fig. 5). We again compared the *k*-mer count correlations of all node pairs within the component (Supplementary Fig. S), distinguishing between the sets of contigs mapping to chromosomes 1 and 8. The former contained 563 nodes, covering 110.32 Mb, of which 315 featured unique *k*-mers and were thus suitable for the correlation computations (covered sequence = 102.48 Mb), whereas the latter contained 527 nodes, covering 74.6 Mb, of which 297 featured unique *k*-mers (covered sequence = 67.05 Mb). Again, we expect to see little or no correlation if two node sets originate from separate chromosomes. In this case, however, the distribution of correlations was consistent with the connections suggested by the assembly graph – contradicting the structure of the Solyntus reference (Supplementary Fig. 5a). These results suggest there is either a large rearrangement that distinguishes between the Altus and Solyntus genomes, or a structural error in the Solyntus reference genome.

## DISCUSSION

We have developed a *de novo* assembly approach that uses accurate long reads and low-depth sequencing data from offspring samples to produce a phased assembly with haplotig lengths up to the length of chromosome arms. To achieve this, our method features multiple innovations. In particular, we designed a complete pipeline that uses haplotype-unique *k*-mers to chromosome sort and phase an assembly graph representing an autopolyploid genome. Importantly, this avoids intermediate steps that flatten the assemblies into contigs, instead resolving the haplotypes directly in the context of the graph topology, which might allow the unified integration of additional data types in the future.

The pseudo-chromosomes resulting from our assembly mapped well to the current monoploid reference genome, but we obtained ∼3.8 times as much sequence data, which indicates comprehensive haplotype resolution. By using low-pass offspring sequencing, our approach is immediately accessible in breeding and research settings where a population of offspring and standard sequencing facilities are available. It avoids the need for single-cell pollen sequencing technology, which is an alternative route to assemblies of comparable quality (Sun et al. 2022).

Despite the rapid advances in phased plant genome assembly, haplotype-resolved chromosome-level assemblies remain challenging for complex autopolyploid genomes. The complete resolution of a haploid human genome foreshadows this development and highlights the methodological advantage of working directly on assembly graphs (Nurk et al. 2021). To resolve the most recalcitrant genomic loci, ultra-long Oxford Nanopore Technologies (ONT) reads have been aligned to assembly graphs constructed from PacBio HiFi reads (Rautiainen and Marschall 2020). We envision that our approach will be combined with such additional data types in future studies. This is currently hampered by difficulties in the preparation of ultra-long sequencing reads (> 100 kb) for plant genomes, but we anticipate the technical challenges will be overcome in the next few years. In our present HiFi-based graphs, shorter contigs tended to lack unique *k*-mers and 12% of the genome was part of such contigs. Mapping additional sequencing data such as ultra-long ONT reads to the graphs could help to bridge the remaining gaps, allowing the inclusion of further graph nodes in the haplotype sequences.

## ONLINE METHODS

### Dosage estimation in unitigs

For each contig, we computed the average coverage by aligning the HiFi reads with the contigs using minimap2. We only considered positions covering the unique sequence of the contig, meaning that overlaps to both neighbouring contigs (if they exist) were not considered. The average coverage *c*_*i*_ of a contig *i* was then computed as the average read depth over all positions. We also computed the total average coverage *m*. Finally, we estimated the dosage *d*_*i*_ of contig *i* as follows:

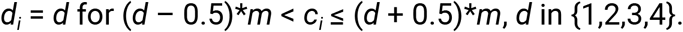

For *c*_*i*_ > 4.5\**m*, we assigned *d*_*i*_ = 5 to denote a repetitive contig.

### Connection of graph components

The clustering of unitigs into haplotype-resolved chromosome clusters involved two steps. First, we attempted to resolve the genome at the chromosome level. Chromosomes may feature several connected components plus additional singletons, so it was necessary to determine which components from the graph belong together. Second, we divided each chromosomal cluster into four distinct clusters, one for each haplotype.

We made use of the previously computed *k*-mer counts in the progeny to cluster unitigs with a similar *k*-mer count pattern. Our clustering procedure followed the idea that we can assign unitigs showing highly similar patterns to the same haplotype, whereas unitigs with opposing patterns are likely to be from the same chromosome but a different haplotype, and unitigs with seemingly unrelated count patterns are probably from different chromosomes.

The similarity between the *k*-mer count patterns of two nodes was assessed by computing the Spearman correlation coefficient (ρ). Two nodes with highly positively correlated *k*-mer count patterns should therefore reflect the same haplotype, whereas highly negative correlations would indicate that the nodes lie on distinct haplotypes. Only nodes from the same chromosome should be highly correlated (positively or negatively), whereas for nodes lying on two separate chromosomes, the *k*-mer counts should be unrelated and any similarities would occur by chance, resulting in low correlation coefficients.

We initially clustered the components and single unitigs into chromosome clusters by grouping all nodes showing the highest pairwise correlation coefficients (ρ > 0.5). These initial clusters were merged when the contigs therein were found to stem from the same graph components. This first clustering step yielded 12 large clusters that were defined as the corresponding chromosomal clusters or pseudo-chromosomes.

### Clustering of unitigs based on similar *k*-mer patterns

To determine the individual haplotypes for each chromosome, we used the previously computed dosage estimation and started by clustering unitigs with dosage 1, because those nodes can only be assigned to a single cluster. We followed an agglomerative method that starts by building seed clusters with the highest correlations and then merges them into larger clusters as well as adding more nodes.

For each node *n* with dosage 1, we initially created one cluster for *n* containing only those unitigs with a high correlation to *n* (ρ > 0.5), producing a set of seed clusters. We then merged these clusters according to the number of common nodes they contain. To distinguish the different linkage groups, we made use of the high negative correlation between two nodes representing different haplotypes.

We created a negative edge between clusters *c*_*i*_ and *c*_*j*_ if there was one node pair (*n*_*i*_, *n*_*j*_), where *n*_*i*_ ∈ *c*_*i*_ and *n*_*j*_ ∈ *c*_*j*_, with high negative correlation (ρ_i,j_ < –0.3). Conversely, we created a positive edge if at least one node pair (*n*_*i*_, *n*_*j*_) existed with a high positive correlation (ρ_i,j_ > 0.5). Two clusters *c*_*i*_ and *c*_*j*_ that were connected by a positive edge could be merged if no contradicting edge existed, such as a positive edge from *c*_*i*_ to another cluster *c*_*k*_ connected negatively to *c*_*j*_.

After the merging steps, all nodes with the highest correlation to other nodes were assigned to clusters. Given that the subset of nodes not highly correlated to any other node (ρ < 0.5) was left out during this procedure, we included these remaining nodes by assigning them to a cluster *c*_*i*_ if the three best hits (the nodes *n*_*a*_, *n*_*b*_ and *n*_*c*_ with the highest correlation to *n*) all belonged to *c*_*i*_. If this was not the case, we were unable to assign *n* unambiguously to a single cluster and it was left unclustered.

Finally, we assigned unitigs with higher dosages to the previously computed haplotype clusters. To cluster a node *n* with dosage *x* (*x* in {2, 3, 4}), we computed the pairwise correlations between *n* and all nodes of all clusters *c*_*1*_, *c*_*2*_, *c*_*3*_ and *c*_*4*_ and added the node to the *x* clusters with the highest ratio of nodes that correlated positively with *n*.

### Assembly of clustered unitigs

Starting with the cluster of contigs for each chromosome, we reconstructed the ordering of contigs throughout the chromosomes to find all possible connections between them in order to create haplotypes with the greatest contiguity. First, we implemented the obvious extensions. If a phased node had only one neighbor in either direction, that neighbor was also considered to be phased. For simple bubble structures (four nodes, including source, sink and two branching nodes) where both the source and the sink node were phased, one of the two branching nodes was assumed to be on the phasing path. If both branches lacked phase, no information was available to pick the correct one, so the node was chosen arbitrarily and the corresponding sequence filled with placeholder characters instead of the node sequence to indicate the absence of correct haplotype sequence information.

We then considered the set of all phased nodes isolated from the rest of the graph. These formed a set of linear block structures, for each of which we were able to identify the two end nodes and recreate the node path (and therefore the sequence) through the block. To also reconstruct the order of these haplotype blocks, we then searched for paths between the end nodes of different blocks that solely contained unphased nodes. For blocks that could be connected uniquely to one additional block, we concatenated the two block sequences and again used placeholder characters for the length of the intervening unphased fragment. Finally, we resolved any remaining overlaps between the extended node paths, resulting in the final output sequences.

## DATA AVAILABILITY

The offspring short reads are available via the NCBI BioProject under accession number PRJEB48582. Similarly, the HiFi reads for Altus are available under accession number PRJNA778192 (Reviewer link https://dataview.ncbi.nlm.nih.gov/object/PRJNA778192?reviewer=cpuvu8qltsolmka5r28nlr9kup).

## CODE AVAILABILITY

The implementation of the workflow described herein is available at the following URL: https://github.com/rebeccaserramari/polyploid-potato-assembly.

## ACKNOWLEDGEMENTS

We acknowledge funding provided by the German Research Foundation (DFG) – 395192176 (to G.W.K. and T.M.) – as well as under Germany’s Excellence Strategy – EXC 2048/1 – 390686111 (to G.W.K. and B.U.), by the German Ministry of Education and Research (BMBF) 031A536C (to B.U.), by the National Institutes of Health (NIH) U01 HG010973 (to T.M.) and by the Dutch TKI top-sector projects *“Genetics Assisted Assembly of Complex Genomes”* (project number BO-68-001-033-WPR) and *“LWV20*.*112 Application of sequence-based multi-allelic markers in genetics and breeding of polyploids”* (project number BO-68-001-042-WPR). We also acknowledge the computational support provided by the Centre for Information and Media Technology (ZIM) at the Heinrich-Heine University Düsseldorf and Dr. Richard M Twyman for manuscript editing.

## AUTHOR CONTRIBUTIONS

R.S.M. and T.M. designed the assembly method, with input from S.S., R.F., B.U and G.W.K. R.S.M. implemented the assembly method, performed the assemblies, and created the figures. R.S.M., S.S., R.F., G.W.K., B.U. and T.M. discussed and interpreted results. R.S.M., R.F., B.U. and T.M. wrote the manuscript, with input from S.S. and G.W.K. P.A. provided offspring sequencing data. M.HW.S. produced HiFi data and ran an initial assembly. All authors read and approved the final manuscript.

## Notes

### Competing Interest Statement

The authors have declared no competing interest.

