## Supplemental Material for "Haplotype-resolved assembly of a tetraploid potato genome using long reads and low-depth offspring data"

**<u>SUPPLEMENTARY MATERIAL</u>**

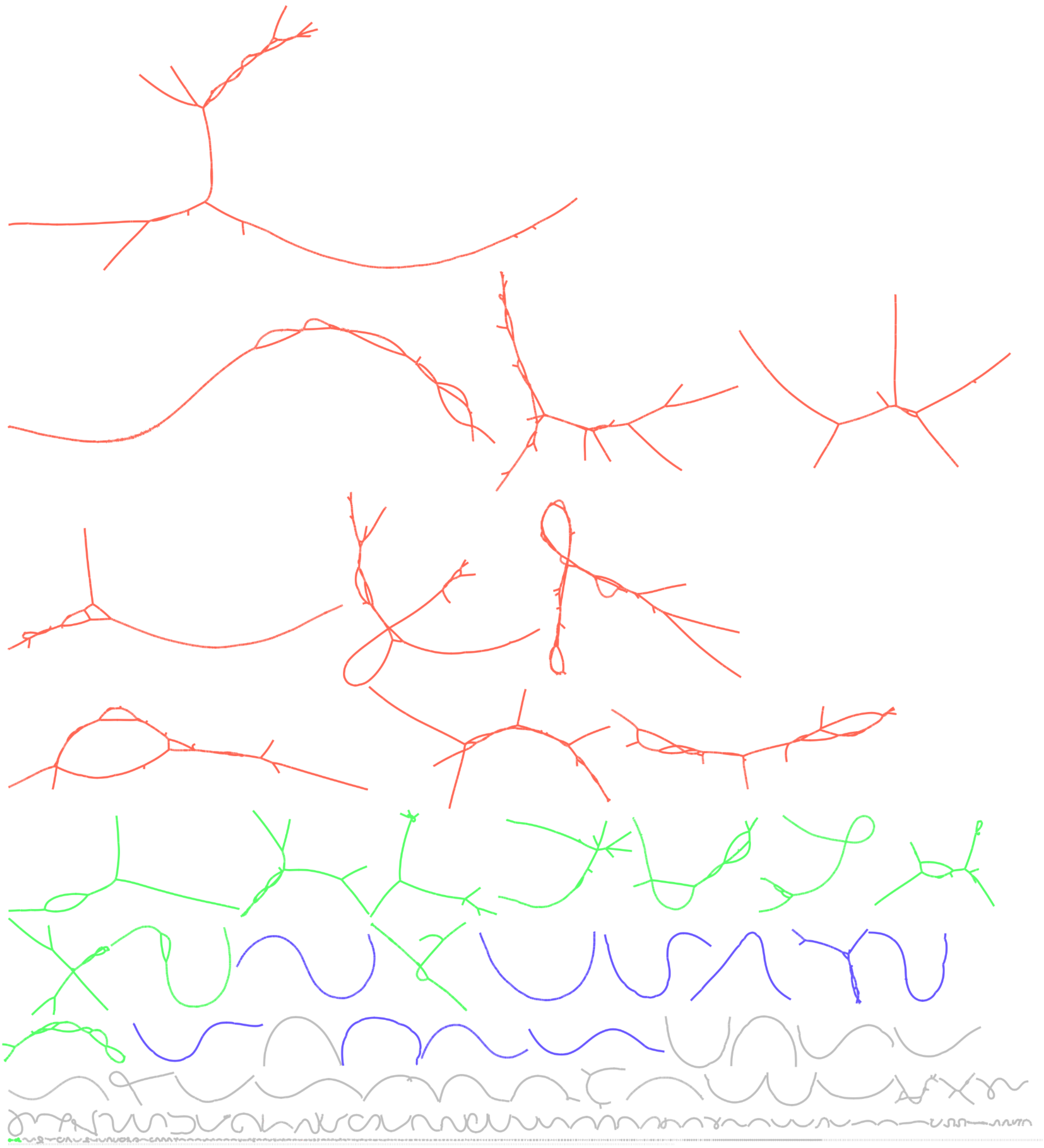

**Suppl. Fig. 1:** Bandage visualisation of the hifiasm raw unitig graph.

The different size categories are indicated by the colouring: red represents the largest components (91–190 Mb), green the second largest (45–66 Mb), and blue the third largest (20–32 Mb).

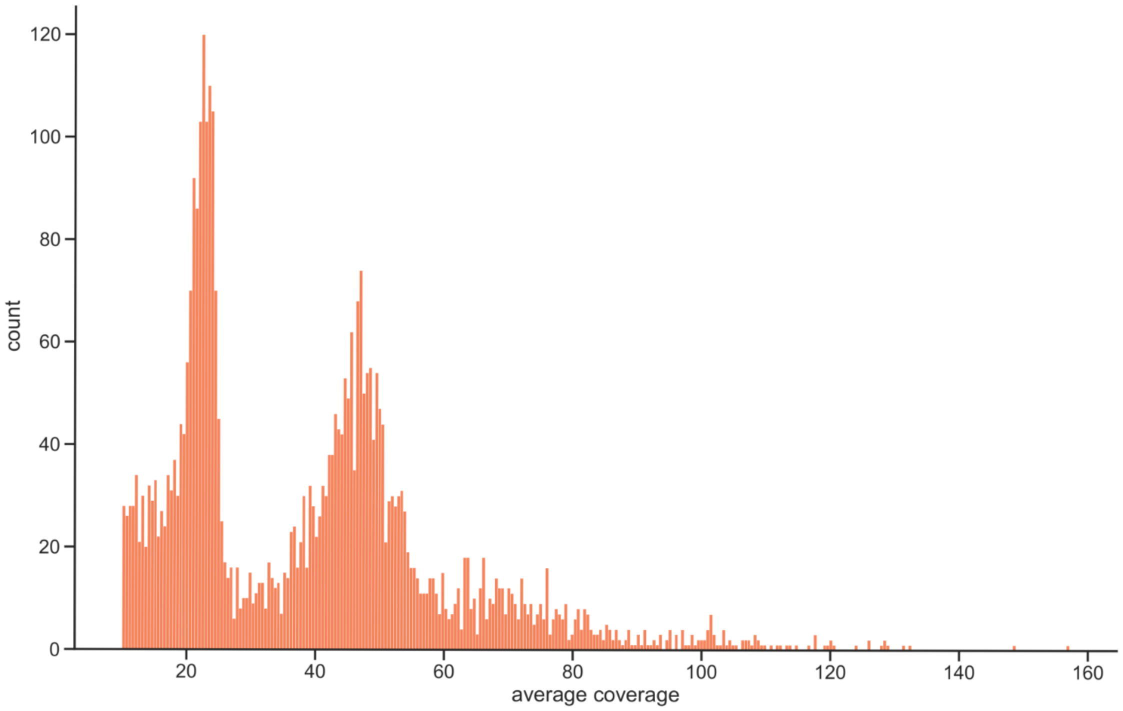


**Suppl. Fig. 2: Dosage distribution of unitigs.**

Contigs with coverage < 10 are filtered out for better visualisation.

**a**

**b**

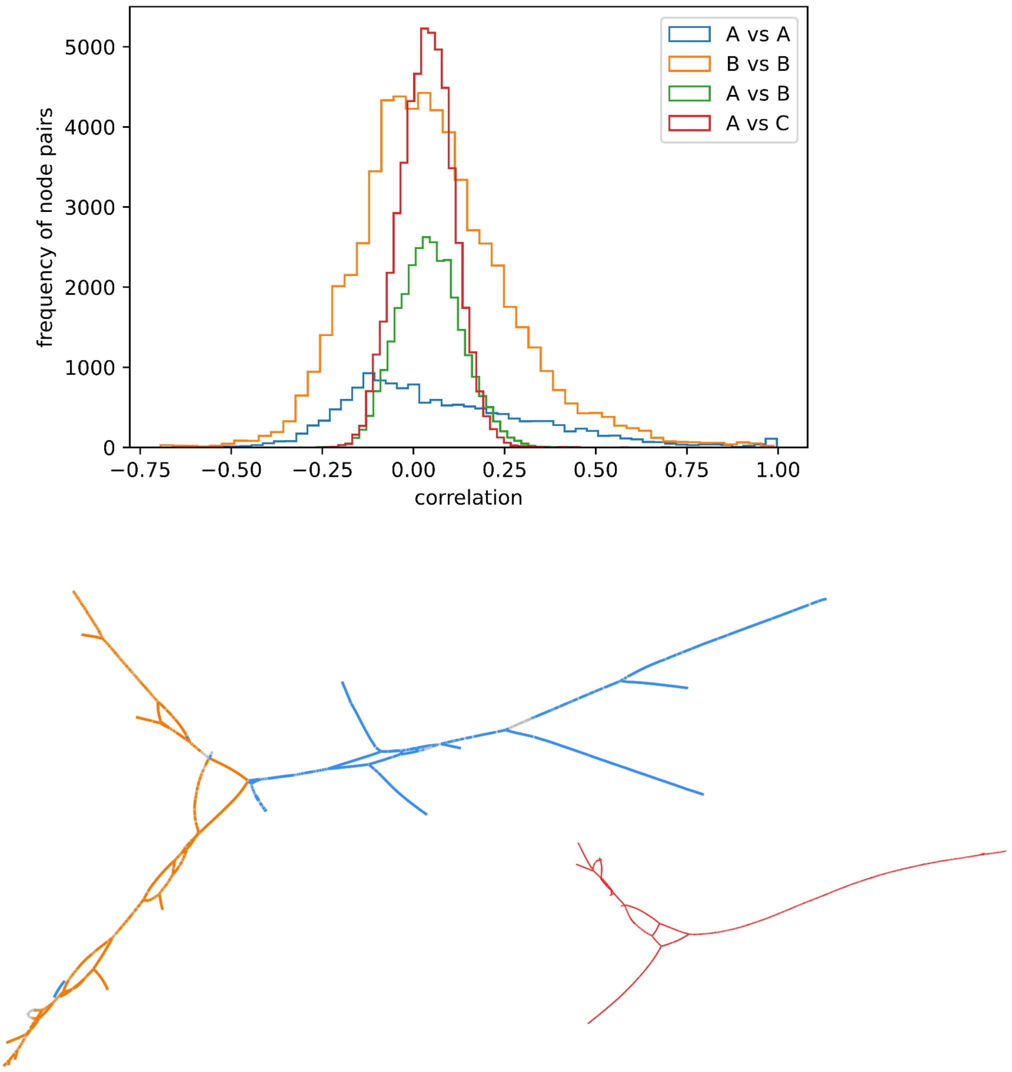


**Suppl. Fig. 3: Analysis of assembly errors in the hifiasm graph between chromosomes 10 and 12 of DMv6.1.**

**a.** Distribution of the pairwise correlation of nodes contained within set A (blue), within set B (orange), between sets A and B (green), and between A and an arbitrarily chosen different component C (red).
**b.** Left: The component of the assembly graph that contains node sets A (contigs that map to chromosome 12 in the DMv6.1 sequence, colored blue) and B (contigs that map to chromosome 10 in the DMv6.1 sequence, colored orange). Right: The component used for comparison, labelled C (contigs are colored in red).

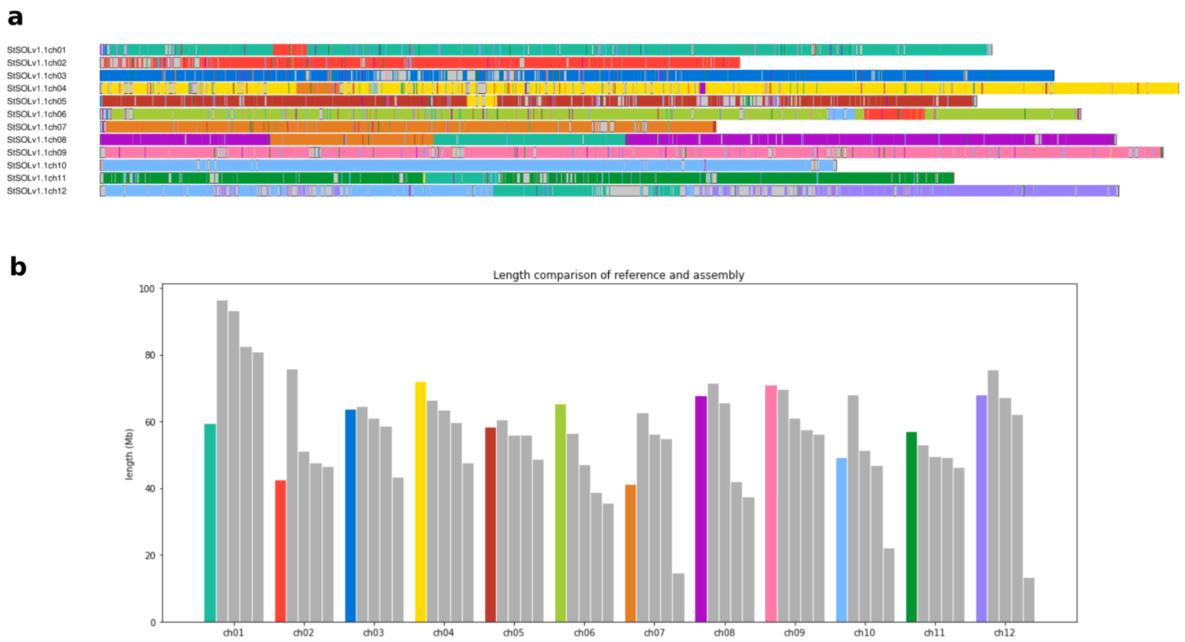


**Suppl. Fig. 4: Mapping of the clusters to the Solyntus v1.1 reference sequence.**

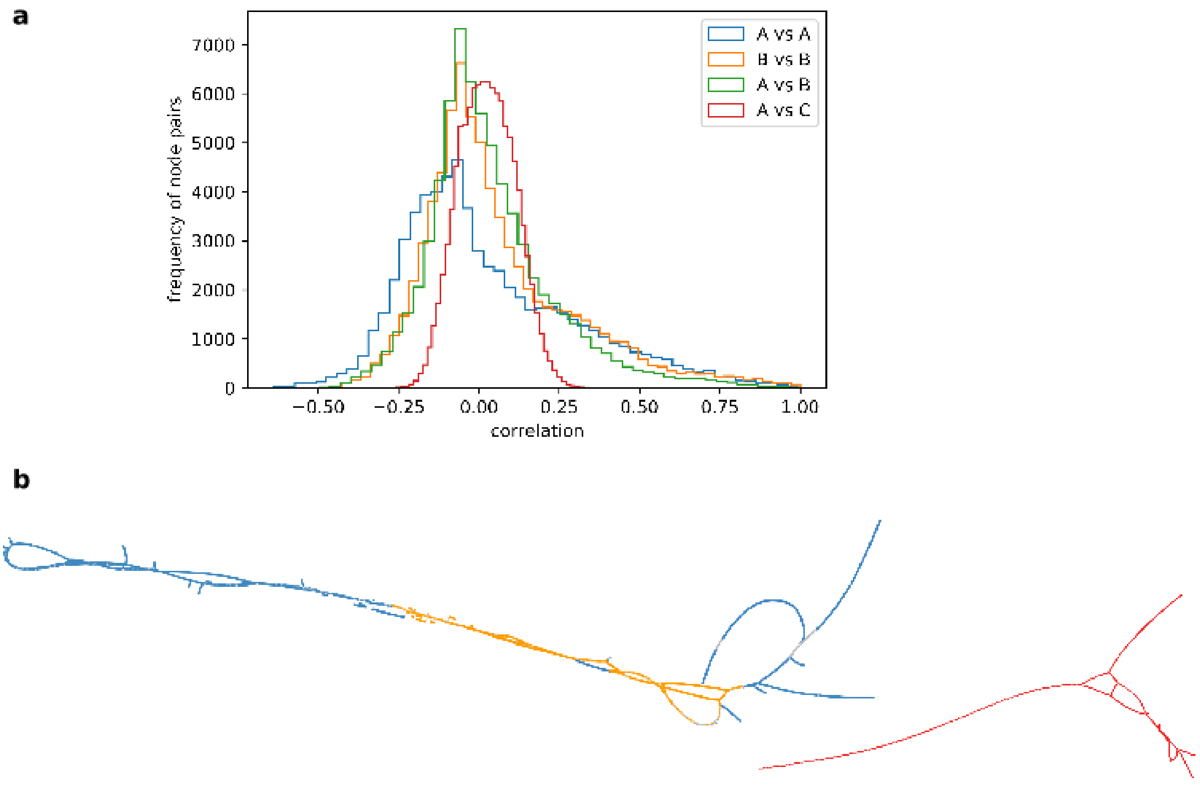


**Suppl. Fig. 5: Analysis of assembly inserts between chromosomes 1 and 8 in the Solyntus v1.1 reference sequence.**

**a.** Distribution of the pairwise correlation of nodes contained within set A (blue), within set B (orange), between sets A and B (green), and between A and an arbitrarily chosen different component C (red).
**b.** Left: The component of the assembly graph that contains node sets A (contigs that map to chromosome 1, colored blue) and B (contigs that map to chromosome 8, colored orange). Right: The component used for comparison, labelled C (contigs are colored in red).
